# DroneMegaDetector, A Generalised Model For Detecting Large Mammals In Aerial Imagery

**DOI:** 10.64898/2026.10.12.758430

**Authors:** Chaim Chai Elchik, Steven Longmore, Dan Morris, Ben Weinstein, Josh Veitch-Michaelis, Camee Knops, Freya Richards, Serge Wich

**Author notes:** Corresponding author: Chaim Chai Elchik.

## Abstract

Large-scale drone surveys can generate imagery across substantial geographic areas, but automated analysis remains challenging because animal appearance, target density, imaging conditions and target scale can vary considerably between surveys. We present a general-purpose RF-DETR Medium object detector developed for large-scale aerial drone imagery and evaluate its ability to generalise across taxa, environments and survey conditions. Unlike specialist counting systems that primarily return point detections, the proposed detector produces explicit bounding boxes, providing a spatial representation that can support counting as well as downstream detection, classification and tracking workflows. We evaluated the detector on an internal held-out test set and independent external datasets spanning cattle, birds, caribou and hippopotamuses, together with imagery acquired under substantially different survey conditions. On the comparative Cattle dataset, RF-DETR matched OWL across the reported point-localisation and counting metrics, while on Caribou it achieved almost identical point-based detection performance and similar aggregate counting error. Performance varied across the broader external benchmark, with reduced transfer to unseen avian imagery and differences between models across datasets. At high altitude on SAVMAP, all evaluated approaches experienced substantial performance degradation, demonstrating a shared limitation associated with small target scale. Overall, the results show that a general-purpose detector trained on heterogeneous drone imagery can provide competitive localisation and counting performance across diverse datasets while retaining explicit bounding-box outputs. The trained model weights and inference code are provided online to support reproducible large-scale aerial wildlife monitoring and further development of drone-based computer-vision workflows.

## 1 Introduction

Drones have become a valuable tool for wildlife monitoring and conservation, offering a way to survey animal populations that is safer, cheaper, and more repeatable than traditional crewed aerial surveys or ground-based counts (Linchant et al. 2015; Tuia et al. 2022; Wich and Koh 2018). By collecting high-resolution imagery over large and often inaccessible areas, drones allow researchers and managers to generate systematic, georeferenced records of species distribution and abundance with a lower risk of observer injury and a lower per-survey cost than manned aircraft (Linchant et al. 2015; Axford et al. 2024). This capability has driven a rapid expansion of drone-based monitoring across taxa and habitats, from savanna ungulates and elephants to marine mammals, birds, and arboreal species, and has positioned drones as a core tool for conservation applications such as population estimation, anti-poaching patrols, and habitat assessment (Aliane 2025; Elchik, Hudson, et al. 2026; Kline, Afridi, et al. 2025; Deng et al. 2026; Lundquist et al. 2025; Allin, Seydou, Frans, et al. 2026).

This growth in data collection has, however, created a new bottleneck: the manual annotation and interpretation of the resulting imagery. Drone surveys can produce thousands of images per flight, and identifying and counting animals in this imagery by hand is slow, labour-intensive, and prone to inconsistency between observers (J. A. J. Eikelboom et al. 2019; Samiappan, B. Santhana Krishnan, et al. 2024b). Even where deep learning is used to accelerate detection, developing an accurate model still depends on large volumes of annotated training data, and the annotation itself is a substantial cost: drawing bounding boxes around individual animals has been reported to take up to seven times longer and cost roughly three times as much as marking a single point on each animal (Ge et al. 2023; Mullen, Tanner, and Sallee 2019). Reducing this annotation and interpretation burden, while retaining the accuracy needed for reliable population estimates, remains a central challenge for scaling drone-based monitoring (Delplanque, Lamprey, et al. 2023).

In response, a large number of deep learning models have been developed to automatically detect animals in drone imagery. The majority of these models, however, have been trained for a specific species, taxonomic group, or study site, including detectors built for dugongs (Maire, Alvarez, and Hodgson 2015), rhinos (Chalmers et al. 2021), dolphins (Bigal et al. 2022), orangutan nests (Teguh et al. 2024; Wich, Ancrenaz, et al. 2025), koalas and other arboreal and thermally distinct species (Backman et al. 2025; Vuong et al. 2025), and elephants (Elchik, Wich, and Burger 2025). While often highly accurate within their intended context, such single-purpose models generally do not generalise well to new species, landscapes, or sensor types, meaning that each new application effectively requires its own bespoke dataset and training effort.

A smaller number of more general, multi-species models have been proposed to address this lack of generalisability. HerdNet, for example, was developed to detect and count multiple African mammal species simultaneously in aerial imagery (Delplanque, Foucher, Théau, et al. 2023; Delplanque, Foucher, Lejeune, et al. 2022), and has since been applied and fine-tuned across several large-scale surveys (Delplanque, Linchant, et al. 2024a; Delplanque, Lamprey, et al. 2023; Allin, Seydou, Radloff, et al. 2026). More recently, the Overhead Wildlife Locator (OWL) benchmarked a weakly supervised, point-based detection approach against HerdNet and other baselines across multiple aerial wildlife datasets (Chacón, Miao, Demuro, C. Robinson, Dodhia, Otarashvili, Holmberg, Larsen, Frederick, Nathan J. Pamperin, et al. 2026b). A common feature of these general models is that they are built around point-level annotations and density estimation rather than bounding boxes, a design choice that substantially reduces annotation cost and effort (Chacón, Miao, Demuro, C. Robinson, Dodhia, Otarashvili, Holmberg, Larsen, Frederick, Nathan J. Pamperin, et al. 2026b; Ge et al. 2023; Mullen, Tanner, and Sallee 2019). This comes at a cost, however: without bounding boxes, these models cannot provide the spatial extent of each detected animal, information that is often needed for downstream tasks such as multi-object tracking, re-identification, or estimating body size and species from imagery. A recent global bird study did, however, use bounding boxes and trained a model for species, age, and sex (Wilson et al. 2026).

To date, no general drone detection model exists that draws bounding boxes for animals in drone imagery, comparable to how MegaDetector filled this role for camera-trap images (Beery, Morris, and S. Yang 2019). The aim of this study is to address this gap by leveraging a large volume of openly available, annotated drone imagery, drawn from across the studies described above and other public sources, to train a general-purpose drone-based animal detector, a “drone megadetector”, that produces bounding-box detections across a wide range of species, habitats, imaging conditions, drone types (as well as one study using a crewed aircraft), sensors, camera angles, and flying heights.

## 2 Methodology

### 2.1 Dataset Composition and Preprocessing

To create the training and validation datasets, we combined proprietary field data with 18 publicly available online datasets (Shao et al. 2020; Adrien Lebreton and Louise Helary 2026; L. Helary and A. Lebreton 2024; Fonod et al. 2025; Samiappan, B Santhana Krishnan, et al. 2024a; sugamg 2026; Delplanque, Linchant, et al. 2024b; Mou et al. 2023; Kline, Kholiavchenko, et al. 2025; Jenna Kline 2025; Kline, Nguyen Ngoc, et al. 2025; Koger et al. 2023; Price, Pranav C Khandelwal, et al. 2025a; Naik et al. 2024; J. A. Eikelboom et al. 2019; Akshatha et al. 2023; Kumar et al. 2024; Praschl et al. 2026), resulting in a total of 19 distinct source datasets. These datasets collectively span four continents (Africa, Asia, Europe, and North America) and capture a highly diverse range of habitats, including wildlife reserves, research centres, and agricultural farms. In total, the aggregated dataset reflects 44 distinct animal taxa, ranging from large African ungulates and elephants to European livestock and Asian primates. A comprehensive breakdown of the geographical locations and specific taxa represented across these 19 datasets is provided in Table 1. The full licensing details and links to the datasets can be found in the supplementary materials.

**Table 1:** Geographical Locations and Identified Taxa Across the 19 Datasets.

| Region | Locations / Habitats | Represented Animal Taxa |
| --- | --- | --- |
| **Africa** | Ethiopia; Mpala Research Center & Ol Pejeta Conservancy (Kenya); Welgevonden Game Reserve (South Africa) | African Buffalo, African Bush Elephant, African Wild Dog, Blue Wildebeest, Bontebok / Blesbok, Chacma Baboon, Common Eland, Common Ostrich, Gelada Monkey, Giraffe, Grevy’s Zebra, Hippopotamus, Impala, Kob, Kudu, Lion, Olive Baboon, Plains Zebra, Red Hartebeest, Thomson’s Gazelle, Topi, Warthog, Waterbuck, White Rhinoceros, Crocodile |
| **Asia** | Cambodia; Agricultural Farms (Japan); China | Amur Tiger, Bactrian Camel, Banteng, Eld’s Deer, Gaur, Giant Panda, Golden snub-nosed Monkey, Lesser Adjutant, Pileated Gibbon, Sambar Deer, Sichuan Takin, Water Buffalo, Kiang |
| **Europe** | Diverse Forests (Austria); Asten (Netherlands); Knowsley Safari & Slapton (United Kingdom); Farms; Coastal Regions | Alpine Ibex, Domestic Cattle, Domestic Dog, Domestic Sheep, Red Deer, Domestic Horse, Seal |
| **North America** | The Wilds (Ohio, USA) | Persian Onager (alongside captive African species) |

All source datasets were first standardised by converting them from their native structures into the Common Objects in Context (COCO) format. During this conversion, specific or varied labels across different datasets (such as specific animal species or vehicle types) were uniformly mapped to three overarching categories: “person” (ID 0), “animal” (ID 1), and “vehicle” (ID 3). Extraneous annotations, such as those representing shadows, were explicitly removed during this phase.

To establish a rigorous evaluation framework and avoid the pitfalls of spatial and temporal data leakage common in ecological machine learning, we structured our data into four distinct splits: training, validation, an internal test set (*test-internal*), and an independent external test suite (*test-external*).

For the 19 primary datasets, all samples were aggregated, randomly reshuffled, and subsequently split into training (70%), validation (20%), and *test-internal* (10%) sets. We explicitly acknowledge that random splitting inherently risks data leakage across geographically or temporally correlated frames; consequently, the *test-internal* split functions primarily as a baseline sanity check for the validation split rather than a definitive measure of real-world performance.

To serve as the true measure of out-of-distribution generalisation and structural robustness, we curated the *test-external* suite. This suite comprises independent datasets entirely held out from the primary training pipeline, carefully selected to probe distinct performance boundaries. It includes data closely mirroring our training characteristics (AED)(Naude and Joubert 2019), high-altitude imagery for evaluating scale invariance (SAVMAP) (Reinhard et al. 2015), out-of-distribution taxonomic data focusing on avian species (Kabra Birds) (Kabra et al. 2022) and unseen large-bodies mammals in the form of Hippopotamus which are also often partially submerged, and the Caribou dataset (Chacón, Miao, Demuro, C. Robinson, Dodhia, Otarashvili, Holmberg, Larsen, Frederick, Nathan J Pamperin, et al. 2026a), which serves as a standard external benchmark. Furthermore, a specific subset of the Shao Cattle dataset (Shao et al. 2020) was designated as a comparative control within this external suite, as it overlaps with the training corpora of both our model and comparison baselines.

To address datasets featuring predominantly small objects, a tiling procedure to 1280×1280 (the intended model training size) was implemented. Datasets were tiled if the average relative bounding box area fell below a 0.00015 coverage threshold. This criterion was established because downscaling such images to a standard training resolution (e.g., 1280 pixels) shrinks the bounding boxes to an absolute pixel size that cannot be resolved by modern architectures like YOLO26 (Sapkota et al. 2025), RF-DETR (I. Robinson et al. 2025), and Grounding DINO (Liu et al. 2024). These models inherently rely on spatial downsampling mechanisms that cause excessively small bounding boxes to effectively disappear, losing the detail necessary for training. In YOLO26, objects whose spatial extent shrinks below the minimum feature-pyramid stride (e.g., *s_min_* = 8) during feature map discretisation fail to encompass any valid anchor centres; consequently, they receive zero positive label assignments and contribute no gradient signal during training. RF-DETR utilises Vision Transformer backbones with discrete patch sizes (e.g., 16 pixels), where sub-patch objects lose their spatial representation and cannot be accurately localised by decoder queries. Similarly, Grounding DINO employs Swin Transformer backbones that extract hierarchical feature scales starting at 4× or 8× downsampling; tiny objects vanish in these downsampled feature maps before they can be successfully aligned with language tokens in the cross-modality decoder.

Because tiling inherently generates a disproportionate number of background images, a dynamic minimum visibility threshold was utilised to filter out bounding box slivers. This threshold was automatically calculated as one-third of the dataset’s average object dimension, bounded by a strict minimum of 5 pixels. Furthermore, to prevent the model from over-indexing on background features, a strict constraint was applied across all datasets to limit the proportion of empty frames (images devoid of annotations) to a maximum of 10% of the total subset. For tiled datasets, satisfying this 10% constraint necessitated a more aggressive random filtering step compared to non-tiled datasets.

After preprocessing, all individual datasets were merged into a single, unified dataset. Original file names were preserved throughout the pipeline by prefixing them with their source dataset identifier. This ensures full traceability, allowing each sample to be mapped back to its origin if required.

The final merged dataset comprises 272,960 total images, encompassing 2,034,611 annotations and utilising 827 GB of storage. To better characterise the dataset’s complexity, we analysed the distribution of instances and spatial footprints. The aggregated dataset contains an average of 7.45 bounding boxes per image (median: 5.00). The mean relative bounding box area is just 0.3487% of the image frame, with a median of 0.1433%, highlighting a severe skew toward small-object detection typical of aerial ecological surveys. Furthermore, to illustrate the high degree of scale, altitude, and background variation across the ‘person’, ‘animal’, and ‘vehicle’ categories, sample imagery from the aggregated dataset is presented in Figure 1. Detailed distributions of image subsets, class labels, and storage requirements are summarised in Tables 2 and 3.

**Figure 1:**
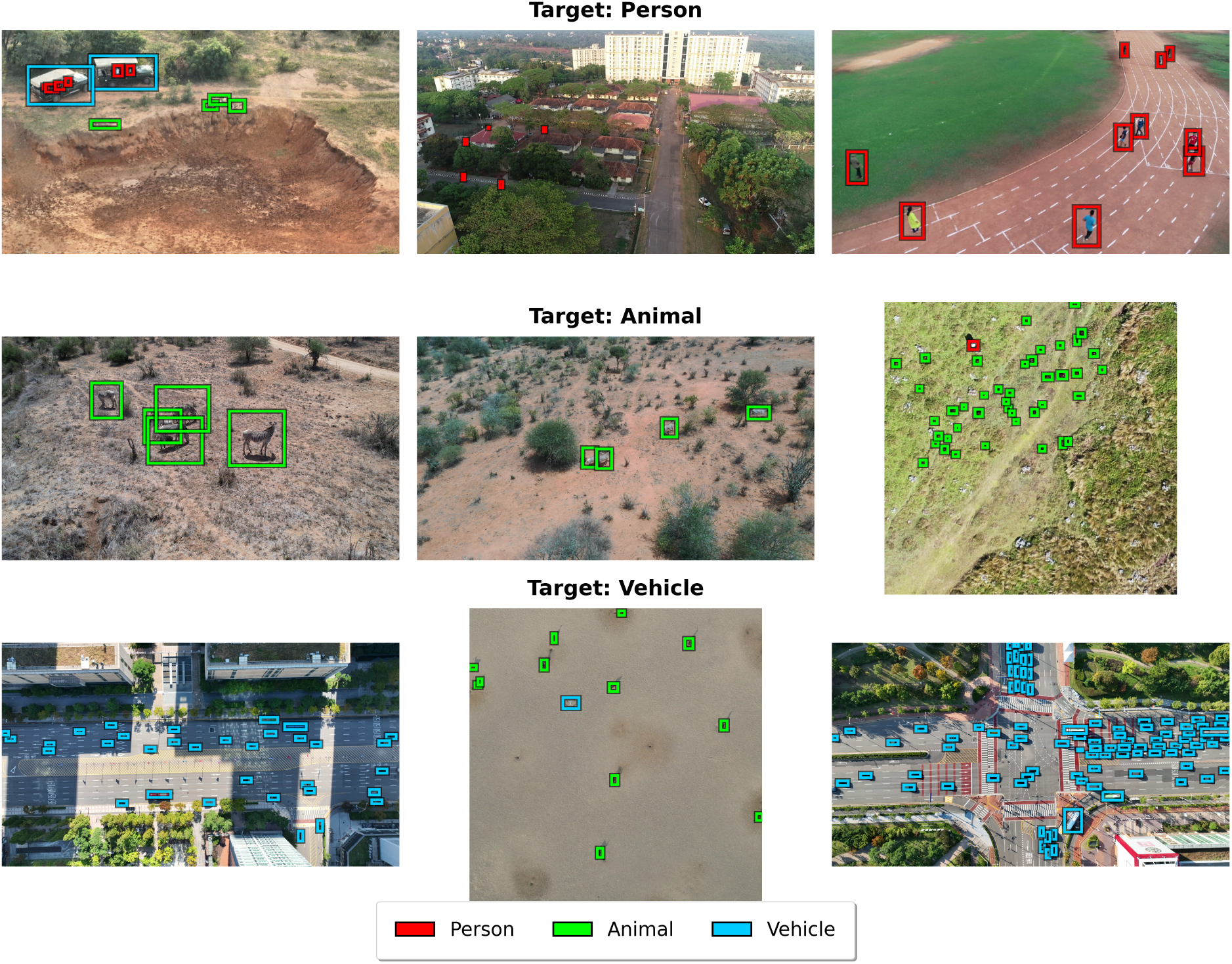
Sample imagery from the aggregated dataset illustrating the high degree of scale and background variation across the target categories.

**Table 2:**
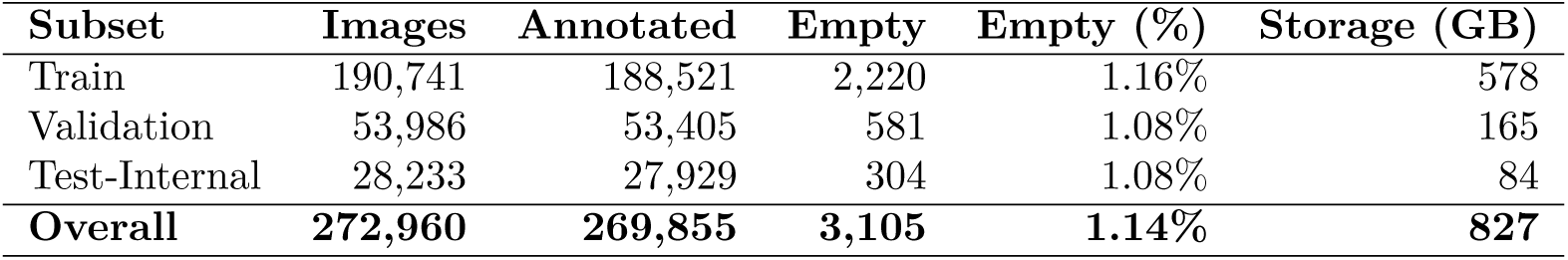
Summary of the aggregated dataset across internal splits.

| Subset | Images | Annotated | Empty | Empty (%) | Storage (GB) |
| --- | --- | --- | --- | --- | --- |
| Train | 190,741 | 188,521 | 2,220 | 1.16% | 578 |
| Validation | 53,986 | 53,405 | 581 | 1.08% | 165 |
| Test-Internal | 28,233 | 27,929 | 304 | 1.08% | 84 |
| **Overall** | **272,960** | **269,855** | **3,105** | **1.14%** | **827** |

**Table 3:** Distribution of annotations across target categories in the aggregated dataset.

| Category | Annotations | Proportion (%) |
| --- | --- | --- |
| Person | 154,692 | 7.60% |
| Animal | 1,606,527 | 78.96% |
| Vehicle | 273,392 | 13.44% |
| **Total** | **2,034,611** | **100.00%** |

### 2.2 Model Selection

We evaluated three distinct object detection architectures to comprehensively benchmark performance across different paradigms: YOLO26, RF-DETR, and Grounding DINO. The YOLO (You Only Look Once) architecture was selected due to its widespread adoption in the computer vision community and its historical precedent in environmental monitoring. Specifically, we utilised YOLO26, the most recent iteration in the YOLO family, to represent the current state-of-the-art in convolutional real-time detectors. To contrast this convolutional approach, we selected RF-DETR, which represents the latest state-of-the-art in image transformer-based real-time detection architectures. Finally, to evaluate generalised vision-language capabilities, we selected Grounding DINO. Grounding DINO is a highly influential open-set detector and also commonly used for fine-tuning applications.

To establish robust baselines and evaluate architectural efficacy, all three models were initially trained on a randomly selected 10% subset of the full unified training dataset. Training was conducted until model convergence, based on no metric improvements on the validation set, using the default hyperparameters to ensure fairness, for each architecture on the Lancaster University High End Computing (HEC) GPU cluster. To ensure parity in hardware acceleration, each training job was allocated 3 NVIDIA V100 32GB GPUs, 30 CPU cores, and 180 GB of memory, running under a maximum 48-hour time limit. Training metrics for all experiments were tracked and logged via Weights & Biases (W&B).

For the YOLO26 implementation, we utilised the medium pretrained weights (yolo26m.pt). Training was conducted with an input image size of 1280 pixels and a total batch size of 12 distributed across 3 GPUs (equating to 4 per GPU). The model was configured to train for a maximum of 300 epochs, featuring a 3-epoch warmup period and an early stopping patience of 15 epochs to guarantee optimal convergence without overfitting. To ensure robust feature learning, a comprehensive suite of drone-specific data augmentations was applied: 180-degree rotations, 10% translation, 50% scaling, 2.0 shear, and 50% probabilities for both vertical and horizontal flips. Additionally, HSV color space augmentations (Hue: 0.015, Saturation: 0.7, Value: 0.4) and a 10% copy-paste augmentation probability were incorporated into the training pipeline.

The RF-DETR model was initialised using the `RFDETRMedium` architecture, similarly configured to process images at a high resolution of 1280 pixels. This transformer-based model was trained for 150 epochs using a base learning rate of 1e-4. Due to the memory constraints inherent to transformer architectures processing high-resolution imagery—which mirrored the constraints of the medium YOLO model-the batch size was similarly restricted to 4 per GPU. To simulate a larger effective batch size and stabilise weight updates, 4 gradient accumulation steps were utilised. During the distributed initialisation phase, a deliberate 30-second synchronisation pause was implemented on secondary GPUs to prevent race conditions during the downloading of the pretrained weights.

Finally, Grounding DINO was fine-tuned using the Swin-Transformer Tiny (Swin-T) backbone, initialised with the `groundingdino_swint_ogc.pth` pretrained weights. The model was distributed across the 3 GPU nodes using the PyTorch distributed run framework (`torchrun`) and relied on standard object detection configuration settings (`cfg_odvg.py`) mapped to our specific dataset parameters.

Following the convergence of the subset models, we evaluated their performance using standard mean average precision (mAP) and F1 scores to determine the optimal architecture for full dataset training. The comparative performance on the 10% subset is summarised in Table 4.

**Table 4:** Performance Evaluation of Candidate Architectures on the 10% Data Subset. F1 scores are evaluated at a standard confidence threshold of 0.5 and an IoU threshold of 0.5.

| Model | Class | F1 Score | mAP@50 | mAP@50-95 |
| --- | --- | --- | --- | --- |
| YOLO26 | Person | 0.7244 | 0.7204 | 0.2658 |
|  | Animal | 0.9412 | 0.9600 | 0.6471 |
|  | Vehicle | 0.9725 | 0.9480 | 0.7275 |
| Grounding DINO | Person | 0.7600 | 0.7630 | 0.2930 |
|  | Animal | 0.9200 | 0.9370 | 0.5240 |
|  | Vehicle | 0.9500 | 0.9460 | 0.6990 |
| RF-DETR | Person | 0.7757 | 0.7542 | 0.2838 |
|  | Animal | 0.9639 | 0.9681 | 0.6604 |
|  | Vehicle | 0.9797 | 0.9576 | 0.7389 |

The evaluation revealed distinct performance variations across the target categories. Grounding DINO achieved the highest mAP@50 (0.7630) for the ‘person’ class, but exhibited lower mAP@50-95 scores across ‘animal’ and ‘vehicle’ targets compared to the other detectors. Conversely, YOLO26 exhibited competitive localisation for the ‘animal’ and ‘vehicle’ classes but struggled significantly with the ‘person’ category, yielding the lowest F1 score (0.7244). RF-DETR demonstrated the most robust and balanced performance across the entire dataset. It achieved the highest overall F1 score (0.7757) for the challenging ‘person’ class, and strictly outperformed both YOLO26 and Grounding DINO in the ‘animal’ and ‘vehicle’ categories, securing the highest stringent mAP@50-95 metrics (0.6604 and 0.7389). Given its balanced accuracy across all ecological and anthropogenic classes, RF-DETR was selected as the optimal architecture to proceed with full-scale dataset training.

### 2.3 Full Model Training

Following the subset evaluation, the RF-DETR architecture was selected for full-scale training on the entirety of the unified dataset. To accommodate the substantially larger volume of data and the intensive computational demands of the transformer architecture, the training pipeline was migrated to the ‘Prospero’ GPU cluster at Liverpool John Moores University using NVIDIA H200 GPUs.

The `RFDETRMedium` model was initialised to process images at a resolution of 1280 pixels. To stabilise the learning process from the outset, an Exponential Moving Average (EMA) mechanism was applied with a decay rate of 0.993, updated at every training step. This temporal smoothing maintains a moving average of the model parameters, effectively acting as a regularizer that reduces weight variance and consistently yields a more stable, robust model with superior generalisation capabilities on unseen data.

Training was conducted using 12 persistent workers per task, with the cuDNN Scaled Dot Product Attention (SDPA) backend explicitly disabled (`torch.backends.cuda.enable_cudnn_sdp(False)`) to ensure strict hardware compatibility. To alleviate the heavy computational burden of the high-resolution transformer architecture and significantly accelerate training time, Automatic Mixed Precision (AMP) was utilised. Empirical monitoring confirmed that this reduction in training precision did not degrade model convergence or final performance. Under this configuration, the model was trained with a batch size of 8 per GPU and no gradient accumulation (step size of 1).

The base learning rate was set to 2 × 10^−4^. Rather than following a linear batch-size scaling rule, this adjustment was made following the official RF-DETR fine-tuning guidelines, which recommend a higher learning rate (e.g., 2 × 10^−4^) for large-scale datasets exceeding 10,000 images to prevent underfitting.

Following these same guidelines, the training schedule was capped at a maximum of 30 epochs. This restricted optimisation horizon was deliberately chosen because extended fine-tuning without the strong “architecture augmentation” regularisation used during the model’s initial Neural Architecture Search (NAS) inherently leads to overfitting (I. Robinson et al. 2025). Furthermore, shorter training schedules are critical to preserving the internet-scale foundational knowledge embedded within the pre-trained DINOv2 backbone, preventing catastrophic forgetting while effectively adapting the model to the target domain.

The full training process required approximately 115.5 wall-clock hours. Training converged at epoch 26, governed by an early stopping mechanism configured with a patience of 10 epochs and a minimum delta of 0.001. For all subsequent evaluations and deployment, the EMA weights of the model were explicitly selected over the standard active training weights.

### 2.4 Evaluation

By evaluating against the *test-external* datasets, we successfully probed distinct performance boundaries: core characteristic generalisation (AED) (Naude and Joubert 2019), scale invariance at high altitudes (SAVMAP) (Reinhard et al. 2015), taxonomic generalisation to unseen avian species (Kabra Birds) (Kabra et al. 2022), and baseline external generalisation (Caribou) (Chacón, Miao, Demuro, C. Robinson, Dodhia, Otarashvili, Holmberg, Larsen, Frederick, Nathan J Pamperin, et al. 2026a). Additionally, the Hippopotamus dataset provided a further test of taxonomic transfer to an unseen large-bodied mammal and consisted of natively non-tiled imagery. Furthermore, the standardised Shao Cattle dataset (Shao et al. 2020) was evaluated as a comparative control. Because this dataset represents an empirically less challenging detection scenario that overlaps with the training corpora of both our model and Microsoft MegaDetector-Overhead (OWL) (Chacón, Miao, Demuro, C. Robinson, Dodhia, Otarashvili, Holmberg, Larsen, Frederick, Nathan J Pamperin, et al. 2026a), it serves to establish a comparative baseline for evaluating model parity on shared data.

This benchmarking framework enabled a direct comparison of the final selected model against two prevailing state-of-the-art baselines: CountGD (Amini-Naieni, Han, and Zisserman 2024) and OWL (Chacón, Miao, Demuro, C. Robinson, Dodhia, Otarashvili, Holmberg, Larsen, Frederick, Nathan J Pamperin, et al. 2026a). Model selection and external generalisation were treated as separate evaluation stages. The in-distribution evaluation was used to determine the model configuration carried forward to the external benchmark, whereas the *test-external* datasets were reserved for assessing generalisation to independent data and were not used for model selection.

Because MegaDetector-Overhead operates natively as a class-agnostic wildlife counting and point-localisation network rather than a standard object detector, standard bounding-box geometric metrics were not used as the primary basis for comparison in the external evaluation. Instead, point-based spatial matching metrics were used to establish a common evaluation framework across architectures. For bounding-box-driven models, predicted bounding boxes were converted to centroid coordinates prior to external evaluation, allowing their detections to be evaluated using the same point representation as OWL and CountGD.

To preserve spatial detail and evaluate high-resolution imagery without inducing extreme downscaling artefacts, models were subjected to localised patch-based processing utilising specific sliding-window resolutions. The RF-DETR was evaluated using an image-slicing resolution of 1280 × 1280 pixels with a 400-pixel overlap. For CountGD, inference was conducted using an 800 × 800 pixel window with an overlap of 250 pixels and the text prompt “animal”, matching the native configurations established in its parent literature. For MegaDetector-Overhead (OWL), the sliding window was configured with a tile size of 512 × 512 pixels and an absolute overlap of 160 pixels to complement its localised density-stitching and peak-detection pathways.

To ensure performance metrics captured the full precision-recall capability of each architecture without artificial truncation, models were permitted to extract deep prediction pools during inference. RF-DETR and YOLO generated spatial predictions down to a near-zero confidence floor of 0.01. CountGD utilised its architecture-specific confidence threshold of 0.3. Conversely, MegaDetector-Overhead bypassed standard confidence scores, utilising its native Local Maxima Detection Strategy (LMDS) with an adaptive threshold of 0.2 and a negative threshold of 0.1 to extract probability peaks.

For the initial model selection and in-distribution baseline evaluations, standard object-detection evaluation was utilised to compare the candidate RF-DETR and YOLO26 models. The spatial accuracy of predicted bounding boxes (*B_pred_*) relative to ground-truth targets (*B_gt_*) was measured via the intersection over union (IoU) criterion: 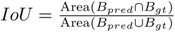 (Ultralytics 2025).

Using an IoU threshold of 0.5, True Positives (*TP*), False Positives (*FP*), and False Negatives (*FN*) were tallied to calculate precision, recall, and F1 scores. Concurrently, standard COCO mean average precision metrics were derived using the pycocotools COCOeval framework. We report mAP@50, corresponding to average precision at an IoU threshold of 0.50, and mAP@50–95, corresponding to the mean AP across IoU thresholds from 0.50 to 0.95 in increments of 0.05. These conventional bounding-box metrics were used for the in-distribution evaluation and model selection because the corresponding training and evaluation annotations provide bounding-box representations of the target animals.

For the point-localisation benchmarking applied to the out-of-distribution data, bounding-box coordinates were converted to their respective centroid coordinates: 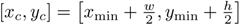.

For point-annotated datasets, the annotated coordinates were used directly as the ground-truth centroids. Spatial matching between predicted points (*p*) and ground-truth centroids (*g*) was determined using a greedy nearest-neighbour matching procedure governed by a spatial distance threshold. Pairwise Euclidean distances were calculated as: 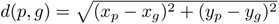.

Predictions were considered sequentially in descending order of confidence. For each prediction, the nearest currently unmatched ground-truth point was selected, provided that its distance was within the applicable spatial tolerance. Each prediction and ground-truth point could therefore be matched at most once. Predictions for which no eligible ground-truth point remained within the tolerance were classified as False Positives, while unmatched ground-truth locations were classified as False Negatives.

Two complementary point-based Average Precision (AP) metrics were calculated. The first, *AP*_40*px*_, uses a fixed 40-pixel spatial tolerance for all ground-truth instances. A prediction is therefore considered a True Positive when *d*(*p, g*) ≤ 40.

This fixed-tolerance AP provides a common spatial criterion across the external datasets and is distinct from conventional IoU-based mAP@50 and mAP@50–95. Whereas COCO AP determines matches according to bounding-box overlap, *AP*_40*px*_ determines matches according to the Euclidean distance between predicted and ground-truth animal centres.

The second metric, *AP_τdyn_*, applies a dynamic spatial tolerance based on the local density of ground-truth animals. For each ground-truth instance *i*, the distance to the nearest other ground-truth centre is calculated as *d*_NN*,i*_ = min*_k_*_̸=*i*_ *d_ik_*. The density-derived tolerance is then set to 0.3*d*_NN*,i*_ and constrained to a minimum of 10 pixels and a maximum of 40 pixels: *τ*_dyn*,i*_ = max [10, min (40,, 0.3*d*_NN*,i*_)].

For point-based datasets, the implementation assumes an effective ground-truth size tolerance of 40 pixels, corresponding to the internal 80 × 80 pixel dummy boxes used to represent point annotations. Consequently, *AP_τdyn_* is density-adaptive for these datasets, with the permissible localisation error decreasing in more densely populated regions while remaining bounded between 10 and 40 pixels. This allows the metric to impose a stricter localisation requirement where neighbouring animals are sufficiently close that a larger tolerance could produce ambiguous assignments.

For both point-based AP metrics, predictions were ranked globally by confidence following image-level matching, and the resulting precision-recall curve was used to calculate Average Precision. AP was calculated using an interpolated precision envelope as *AP* = ∑*_n_*(*R_n_ − R_n_*_−1_)*P_n_*, where *R_n_* and *P_n_* represent recall and precision at successive confidence-ranked operating points. Consequently, *AP*_40*px*_ and *AP_τdyn_* are threshold-independent with respect to the final reported confidence operating point: they evaluate the complete confidence ranking rather than performance at one arbitrarily selected confidence threshold.

Additional operating-point statistics were extracted from the same confidence-ranked predictions. Precision and recall at specified target recall levels were calculated by identifying the first confidence-ranked operating point at which recall reached or exceeded the target value. We report precision at 95%, 85%, 75%, 65%, and 50% recall as P@R95, P@R85, P@R75, P@R65, and P@R50, respectively. Where the target recall was not reached by the available predictions, the corresponding precision was recorded as zero. These statistics complement AP by describing the precision achieved at specific points along the confidence-ranked precision-recall curve.

For environmental monitoring, absolute counting accuracy across large spatial extents frequently supersedes exact pixel-level localisation. We therefore quantified absolute count discrepancies via Mean Absolute Error (MAE) and Root Mean Square Error (RMSE). Because the confidence-score distributions and native inference thresholds differ between architectures, the counting threshold was optimised independently for each model rather than imposing a common fixed confidence threshold. Candidate thresholds were obtained from the unique prediction confidence scores, with up to 101 percentile-derived candidates used when more than 100 scores were available. The threshold *t*^∗^ was selected as the value minimising mean absolute counting error across the evaluation images.

For each image *i*, the counting error was calculated as 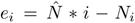, where 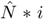 is the number of predictions exceeding *t*^∗^ and *N_i_* is the ground-truth count. MAE and RMSE were then calculated as 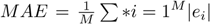, and 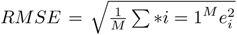, where *M* is the number of evaluated images. MAE therefore represents the mean absolute image-level counting error, while RMSE places greater weight on larger image-level counting errors. Ninety-five percent confidence intervals for both metrics were obtained using bootstrap resampling of the image-level errors (*B* = 1, 000).

Finally, aggregate counting bias was quantified using the total predicted and ground-truth counts and their relative difference: 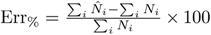. The corresponding Pred/GT values report the total number of predicted and ground-truth animals at the model-specific counting threshold *t*^∗^. Positive Err% values indicate aggregate overcounting, whereas negative values indicate aggregate undercounting. Unlike MAE and RMSE, which quantify the magnitude of image-level counting errors, Err% captures the net deviation in the total estimated population.

This evaluation design therefore separates conventional in-distribution object-detection performance used for model selection from point-based external generalisation and counting performance. In-distribution model selection was based on conventional COCO bounding-box metrics, including mAP@50 and mAP@50–95. The independent external benchmark instead used centre-distance-based *AP*_40*px*_ and *AP_τdyn_*, supplemented by P@R95, P@R85, P@R75, P@R65, and P@R50, to evaluate individual-animal detection and localisation across heterogeneous annotation formats and architectures. Counting performance was evaluated separately using MAE, RMSE, Pred/GT, and aggregate Err%, with the counting confidence threshold independently optimised for each model.

### 2.5 Code and model availability

The inference code and trained RF-DETR Medium model weights will be made available via GitHub at https://github.com/ConservationDronesAI/DroneMegaDetector and at HuggingFace via https://huggingface.co/ConservationDrones/DroneMegaDetector. The inference scripts supports both single-pass and tiled processing of input images, with user-configurable confidence thresholds and output paths for annotated detections. Tiled inference is implemented using 1280 × 1280 image slices, while single-pass inference processes the complete input image. The script automatically selects an available computational device, prioritising Apple Silicon Metal Performance Shaders (MPS), followed by CUDA-enabled GPUs and CPU execution. It also reads the native inference resolution from the model checkpoint when available and applies inference optimisation before prediction. The resulting detections are returned as bounding boxes with associated class identifiers and confidence scores, which can be visualised and exported as annotated images. The trained model weights will be provided in the same online repository as the inference code, supporting local deployment and evaluation of the released detector.

## 3 Results

### 3.1 Performance on Internal Held-Out Test Data

The fully trained RF-DETR Medium model was first evaluated on the internal held-out test split to establish a baseline of its in-domain capabilities. The architecture demonstrated exceptional localisation and classification accuracy for the “Animal” and “Vehicle” categories, achieving F1 scores of 0.9717 and 0.9637, respectively. The mean Average Precision (mAP@50) for these classes mirrored this high performance. However, the model exhibited comparative difficulty with the “Person” class, yielding an F1 score of 0.7045. This reduction in accuracy is driven primarily by a lower precision score (0.6785).

**Table 5:** RF-DETR Performance on the Internal Held-Out Test Split (Untiled). Precision and recall represent absolute point estimates calculated at the optimal F1 confidence threshold of 0.5 and a spatial IoU threshold of 0.5.

| Class | Precision | Recall | F1 Score | mAP@50 | mAP@50-95 |
| --- | --- | --- | --- | --- | --- |
| Animal | 0.9694 | 0.9740 | 0.9717 | 0.9694 | 0.7580 |
| Person | 0.6785 | 0.7326 | 0.7045 | 0.6785 | 0.2553 |
| Vehicle | 0.9599 | 0.9676 | 0.9637 | 0.9599 | 0.7866 |

### 3.2 Performance on the Comparative Control Dataset

The standardised Shao Cattle dataset (Shao et al. 2020) provided a comparative control in which the evaluation conditions were shared across the established counting baselines and the proposed detector. Under the point-based evaluation, RF-DETR and OWL achieved identical performance, with *AP*_40*px*_ = 0.991 and *AP_τdyn_* = 0.991. Both models produced 1,854 predictions for 1,836 ground-truth animals, corresponding to an aggregate error of only +0.98%. Their image-level counting errors were also identical, with an MAE of 0.23 animals (95% CI: 0.19–0.28) and an RMSE of 0.63 animals. Precision remained above 0.99 at all reported recall levels from P@R85 onwards and was 0.969 at 95% recall.

CountGD, evaluated on the corresponding Cattle imagery using its tiled configuration, also achieved high point-based performance, with *AP*_40*px*_ = 0.929 and *AP_τdyn_* = 0.929. Its aggregate count was 1,773 animals against 1,836 ground-truth animals (−3.43%), with an MAE of 0.77 and RMSE of 1.70. Precision at 95% recall was 0.659, increasing to 0.998 at 50% recall. Thus, on the comparative control, RF-DETR achieved point-localisation and counting performance equivalent to OWL and substantially reduced the apparent counting gap observed under the previous evaluation.

**Table 6:** Point-based detection and counting performance on the comparative control dataset (Shao Cattle).

| Model | $AP_{40px}$ | $AP_{\tau_{dyn}}$ | P@R95 | P@R85 | P@R50 | MAE | RMSE | Err% |
| --- | --- | --- | --- | --- | --- | --- | --- | --- |
| RF-DETR | 0.9909 | 0.9909 | 0.9694 | 0.9955 | 0.9989 | 0.23 | 0.63 | +0.98 |
| OWL | 0.9909 | 0.9909 | 0.9694 | 0.9955 | 0.9989 | 0.23 | 0.63 | +0.98 |
| CountGD | 0.9293 | 0.9288 | 0.6592 | 0.9091 | 0.9978 | 0.77 | 1.70 | -3.43 |

### 3.3 Generalisation to Out-of-Distribution Taxa and Environments

Across the independent *test-external* datasets, the results provide no evidence of a general counting-performance penalty associated with the proposed detector. Instead, RF-DETR remained competitive with the specialist counting baselines across substantially different taxa and imaging conditions, although the relative performance of the models varied between datasets.

On AED, OWL achieved the highest point-based AP, with *AP*_40*px*_ = 0.899 and *AP_τdyn_* = 0.857, compared with 0.886 and 0.816 for RF-DETR. OWL also produced the lowest counting error, with an MAE of 1.12 animals and an aggregate error of +2.53%. RF-DETR produced 2,634 predictions for 2,970 ground-truth animals, corresponding to a moderate aggregate undercount of −11.31% and an MAE of 1.89. Nevertheless, RF-DETR maintained high precision across the recall range, increasing from 0.518 at P@R95 to 0.949 at P@R50. CountGD performed substantially lower on this dataset, with *AP*_40*px*_ = 0.192 and an aggregate undercount of −67.07%.

The Kabra Birds dataset produced a different pattern. CountGD achieved the highest AP, with *AP*_40*px*_ = 0.887 and *AP_τdyn_* = 0.868, followed by OWL at 0.823 and 0.813. RF-DETR achieved lower localisation AP of 0.657 and 0.644, respectively. However, the corresponding counting results again demonstrate that detection AP and counting accuracy are not interchangeable. OWL produced the lowest image-level counting error among the three models, with an MAE of 13.48 animals and an aggregate error of −2.59%, compared with an MAE of 22.50 and an aggregate error of +3.35% for CountGD. RF-DETR showed a larger undercount of −37.60% and an MAE of 50.52. Despite this counting error, RF-DETR achieved 0.955 precision at P@R50, indicating that its principal limitation on this dataset was recall rather than the precision of detections that were retained.

On the Caribou dataset, all three architectures achieved high point-based detection performance. OWL and RF-DETR produced almost identical *AP*_40*px*_ values of 0.9690 and 0.9694, respectively, while CountGD achieved 0.8530. Under the density-adaptive metric, OWL reached 0.9344, compared with 0.9288 for RF-DETR and 0.7805 for CountGD. The counting results similarly showed close agreement between RF-DETR and OWL: RF-DETR produced 12,621 predictions for 12,456 ground-truth animals (+1.32%), while OWL produced 12,435 (−0.17%). Their MAEs were 0.84 and 0.41 animals, respectively. Both maintained precision above 0.99 at P@R50.

The Hippopotamus dataset provided a further test of transfer to an unseen large-bodied mammal under natively non-tiled imagery. OWL achieved the highest point-based AP, with *AP*_40*px*_ = 0.763 and *AP_τdyn_* = 0.700, followed by RF-DETR at 0.719 and 0.678. CountGD achieved substantially lower values of 0.508 and 0.407. Importantly, the counting results again placed RF-DETR relatively close to OWL: RF-DETR produced an aggregate overcount of +61.86% and an MAE of 1.48, whereas OWL produced an undercount of −9.37% and an MAE of 1.16. The high aggregate error for RF-DETR therefore reflects a systematic threshold-calibration difference rather than a complete failure to localise individual animals, as reflected by its *AP*_40*px*_ of 0.719 and P@R50 of 0.958.

Overall, the external results indicate that RF-DETR does not exhibit the systematic counting deficit suggested by the earlier evaluation. OWL remained stronger on several datasets, particularly AED and Hippopotamus, while CountGD achieved the highest AP on Kabra Birds. However, RF-DETR achieved near-identical point-localisation performance to OWL on Caribou, identical performance on the comparative Cattle control, and competitive performance on the remaining external datasets. The results therefore indicate that the proposed detector can approach the counting and point-localisation performance of specialist wildlife counting systems while additionally producing explicit bounding-box detections.

**Table 7:** Point-based detection and counting performance across the out-of-distribution datasets. P@R95, P@R85, P@R75, P@R65 and P@R50 denote precision at the first confidence-ranked operating point reaching 95%, 85%, 75%, 65% and 50% recall, respectively. MAE and RMSE are in animals per image.

| Dataset | Model | $AP_{40px}$ | $AP_{\tau_{dyn}}$ | P@R95 | P@R85 | P@R75 | P@R65 | P@R50 | MAE | RMSE | Err% |
| --- | --- | --- | --- | --- | --- | --- | --- | --- | --- | --- | --- |
| AED | CountGD | 0.192 | 0.153 | 0.000 | 0.000 | 0.000 | 0.000 | 0.171 | 6.46 | 11.28 | -67.07 |
|  | OWL | 0.899 | 0.857 | 0.761 | 0.897 | 0.918 | 0.928 | 0.937 | 1.12 | 3.09 | +2.53 |
|  | RF-DETR | 0.886 | 0.816 | 0.518 | 0.805 | 0.868 | 0.910 | 0.949 | 1.89 | 3.91 | -11.31 |
| Kabra Birds | CountGD | 0.887 | 0.868 | 0.000 | 0.863 | 0.898 | 0.921 | 0.946 | 22.50 | 53.70 | +3.35 |
|  | OWL | 0.823 | 0.813 | 0.000 | 0.764 | 0.877 | 0.922 | 0.950 | 13.48 | 22.12 | -2.59 |
|  | RF-DETR | 0.657 | 0.644 | 0.000 | 0.000 | 0.000 | 0.624 | 0.955 | 50.52 | 110.66 | -37.60 |
| Caribou | CountGD | 0.853 | 0.781 | 0.000 | 0.937 | 0.970 | 0.979 | 0.981 | 1.03 | 1.83 | -5.66 |
|  | OWL | 0.969 | 0.934 | 0.969 | 0.987 | 0.991 | 0.994 | 0.995 | 0.41 | 0.79 | -0.17 |
|  | RF-DETR | 0.969 | 0.929 | 0.893 | 0.961 | 0.980 | 0.988 | 0.994 | 0.84 | 1.84 | +1.32 |
| Hippopotamus | CountGD | 0.508 | 0.407 | 0.000 | 0.000 | 0.000 | 0.000 | 0.931 | 2.75 | 4.55 | -43.39 |
|  | OWL | 0.763 | 0.700 | 0.000 | 0.485 | 0.725 | 0.854 | 0.931 | 1.16 | 1.78 | -9.37 |
|  | RF-DETR | 0.719 | 0.678 | 0.008 | 0.078 | 0.408 | 0.824 | 0.958 | 1.48 | 2.46 | +61.86 |

### 3.4 Scale Invariance and High-Altitude Limitations

The SAVMAP dataset exposed a substantially different limitation associated with high-altitude imagery and small target size. All three architectures experienced a pronounced reduction in point-based detection performance. RF-DETR achieved *AP*_40*px*_ = 0.126 and *AP_τdyn_* = 0.065, while OWL achieved 0.135 and 0.068. CountGD performed substantially lower, with *AP*_40*px*_ = 0.013 and *AP_τdyn_* = 0.010.

The counting results showed the same degradation. RF-DETR produced 1,962 predictions for 7,474 ground-truth animals, corresponding to an aggregate undercount of −73.75%, while OWL produced 2,268 predictions (−69.65%). Their MAEs were 9.30 and 8.85 animals per image, respectively. CountGD produced only one prediction across the evaluation set, resulting in an aggregate error of −99.99% and an MAE of 11.43. Precision at the reported recall levels was effectively zero for all models, with RF-DETR reaching only P@R50 = 0.022.

Thus, the high-altitude SAVMAP benchmark represents a common performance boundary across all evaluated architectures rather than a limitation specific to the proposed detector. The relatively small difference between RF-DETR and OWL in both *AP*_40*px*_ and *AP_τdyn_*, together with their similarly severe counting errors, indicates that the loss of performance at this scale is shared across the tested detection paradigms.

**Table 8:**
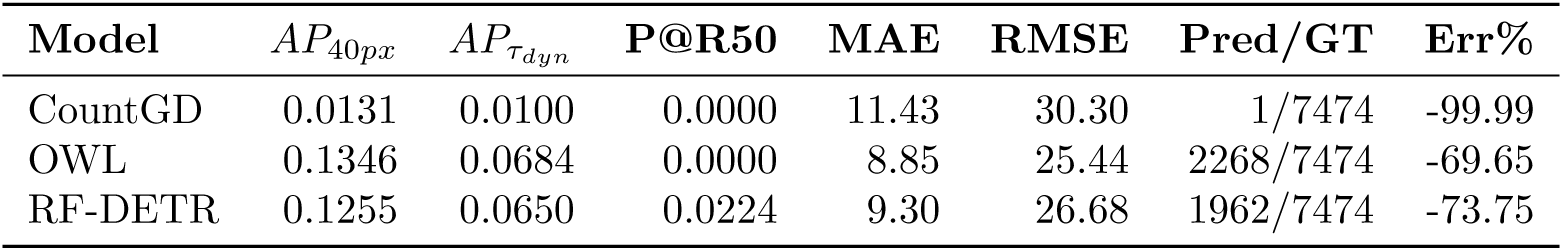
Point-based detection and counting performance on the high-altitude SAVMAP dataset.

| Model | $AP_{40px}$ | $AP_{\tau_{dyn}}$ | P@R50 | MAE | RMSE | Pred/GT | Err% |
| --- | --- | --- | --- | --- | --- | --- | --- |
| CountGD | 0.0131 | 0.0100 | 0.0000 | 11.43 | 30.30 | 1/7474 | -99.99 |
| OWL | 0.1346 | 0.0684 | 0.0000 | 8.85 | 25.44 | 2268/7474 | -69.65 |
| RF-DETR | 0.1255 | 0.0650 | 0.0224 | 9.30 | 26.68 | 1962/7474 | -73.75 |

## 4 Discussion

### 4.1 A general-purpose detector for aerial wildlife imagery

The results demonstrate that a general-purpose detector trained on heterogeneous aerial imagery can retain competitive counting performance while providing explicit bounding-box detections. On the comparative Cattle dataset, RF-DETR matched OWL on both point-localisation and counting metrics, with identical *AP*_40*px*_, *AP_τdyn_*, MAE, RMSE, aggregate count error and precision at the reported recall levels. On the Caribou dataset, RF-DETR and OWL likewise achieved almost identical point-based detection performance, with *AP*_40*px*_ values of 0.969 and 0.969, respectively, and similar aggregate counting errors of +1.32% and −0.17%. These results provide no evidence of a general trade-off in which retaining bounding-box outputs necessarily results in poorer counting performance than specialist point-based systems.

However, the relative performance of the models varied across datasets. OWL retained advantages on AED and Hippopotamus, while CountGD achieved the highest point-based AP on Kabra Birds. RF-DETR therefore cannot be considered uniformly equivalent to the specialist systems across all taxa and imaging conditions. Rather, the external benchmark indicates that a general-purpose bounding-box detector can operate within the same broad performance range as specialist counting systems on several datasets, while retaining richer spatial outputs. This distinction is important because the principal contribution of the proposed detector is not to replace specialised counting systems under every evaluation condition, but to provide a general detection layer that can support counting alongside downstream analyses requiring explicit object localisation.

### 4.2 Generalisation across taxa and survey conditions

The external datasets provide evidence that the proposed detector can transfer to taxa and imaging conditions that differ substantially from its training distribution, although the magnitude of this transfer varies. On AED, RF-DETR achieved *AP*_40*px*_ = 0.886 and *AP_τdyn_* = 0.816, approaching OWL at 0.899 and 0.857. Its counting performance was weaker than OWL, with an MAE of 1.89 compared with 1.12 and an aggregate undercount of −11.31% compared with +2.53%. Nevertheless, RF-DETR retained high precision at lower recall operating points, increasing from 0.518 at P@R95 to 0.949 at P@R50. CountGD performed substantially worse on this dataset, with an *AP*_40*px*_ of only 0.192 and an aggregate undercount of −67.07%. Thus, although RF-DETR did not match OWL on AED, it retained strong point-localisation performance while providing bounding-box outputs.

The Kabra Birds dataset represents a more substantial taxonomic transfer, as avian targets were not represented in the training data. Here, RF-DETR achieved lower localisation AP than both specialist methods, with *AP*_40*px*_ = 0.657 and *AP_τdyn_* = 0.644, compared with 0.887 and 0.868 for CountGD and 0.823 and 0.813 for OWL. Its counting performance was correspondingly weaker, with an aggregate undercount of −37.60% and an MAE of 50.52 animals per image. However, its precision at P@R50 was 0.955, despite reaching only 0.624 at P@R65 and failing to reach the higher reported recall levels. This pattern suggests that reduced recall, rather than poor precision among the detections retained at lower recall, was an important contributor to its lower counting performance on this dataset. The result nevertheless demonstrates that useful detection can transfer to taxa not explicitly represented in the training distribution, while also highlighting the limits of this transfer.

The Caribou dataset showed substantially stronger generalisation. RF-DETR and OWL achieved almost identical *AP*_40*px*_ values of 0.9694 and 0.9690, respectively, while RF-DETR reached *AP_τdyn_* = 0.9288 compared with 0.9344 for OWL. Their aggregate count errors were also close, at +1.32% for RF-DETR and −0.17% for OWL, although OWL achieved a lower MAE of 0.41 compared with 0.84 for RF-DETR. Both models maintained precision above 0.99 at P@R50. These results provide a second example, alongside the Cattle control, in which a general-purpose bounding-box detector approaches the point-based performance of a specialist system without an obvious counting disadvantage.

The Hippopotamus dataset provided a further test of transfer to an unseen large-bodied mammal under natively non-tiled imagery. OWL achieved higher point-based AP than RF-DETR, with *AP*_40*px*_ = 0.763 and *AP_τdyn_* = 0.700, compared with 0.719 and 0.678 for RF-DETR. RF-DETR nevertheless remained substantially above CountGD, which achieved 0.508 and 0.407, respectively. The counting results require more careful interpretation. At the model-specific threshold selected to minimise image-level MAE, RF-DETR produced 13,122 predictions for 8,107 ground-truth animals, corresponding to an aggregate error of +61.86%, while its MAE was 1.48 animals per image. OWL produced 7,347 predictions, corresponding to −9.37% aggregate error and an MAE of 1.16.

The distinction between these metrics is important. The aggregate error measures the net difference between the total number of predictions and ground-truth animals across the dataset, whereas MAE measures the absolute image-level counting error and was itself used to select the RF-DETR operating threshold. RF-DETR’s high aggregate overcount therefore does not imply a uniform 61.86% counting error across images. Instead, it reflects the behaviour of the MAE-optimised threshold across images with very different target densities. RF-DETR produced relatively few high-confidence detections in densely populated images, so lowering the confidence threshold recovered additional low-confidence detections and reduced false-negative counting error in those images. The same threshold, however, admitted many false-positive detections in images containing fewer or no hippos, resulting in a substantial positive aggregate bias. The P@R profile is consistent with this threshold-dependent behaviour: precision was only 0.008 at P@R95 and 0.078 at P@R85, but increased to 0.408 at P@R75, 0.824 at P@R65 and 0.958 at P@R50. Thus, the Hippopotamus results indicate a calibration and operating-point limitation rather than a complete failure of object localisation.

### 4.3 Bounding boxes as both outputs and interfaces

The external evaluation deliberately converted RF-DETR bounding boxes to centroids before comparison with the point-based baselines. This provides a common basis for evaluating localisation and counting while ensuring that differences in output representation do not themselves determine the benchmark outcome. The results indicate that retaining the richer bounding-box representation does not necessarily impose a substantial counting-performance penalty relative to specialist point-based systems.

Bounding boxes nevertheless provide substantially more information than a single point location. They preserve object extent and spatial occupancy, and can therefore act as an interface between detection and subsequent ecological computer-vision tasks. For example, bounding-box detections can be passed to downstream species classification, tracking, behaviour analysis or individual re-identification pipelines. Recent wildlife applications have similarly used object detections as the first stage of more complex analytical workflows, including automated monitoring and behavioural analysis Duporge et al. 2024; Price, Pranav C. Khandelwal, et al. 2025b; Troscianko et al. 2025; Zábó et al. 2026. A general-purpose detector can therefore provide a common detection layer across multiple applications rather than requiring a separate counting-specific model for each downstream task.

This flexibility is particularly relevant for aerial surveys, where the same imagery may support several analytical objectives. A detector capable of identifying animals, people and vehicles while returning spatially explicit bounding boxes can provide a reusable first-stage representation for ecological monitoring and operational screening. Counting can then be performed from the same detections without requiring a separate point-based inference system. The present results suggest that this additional output structure need not inherently compromise counting performance.

The bounding-box representation does, however, remain a two-dimensional image-space representation. Tracking detections across frames can provide trajectories, but conversion to absolute animal movement, body dimensions or three-dimensional positions requires additional calibration, georeferencing or depth information. Consequently, the proposed detector should be considered a general detection component rather than a complete ecological measurement system.

### 4.4 Practical deployment and future development

A general-purpose detector is most useful when it can be deployed across different hardware and survey environments without substantial changes to the inference pipeline. The proposed framework was designed for local inference and can operate using standard computer-vision hardware, including GPU, Apple Silicon and CPU-based systems. The released model weights, inference scripts and associated dependencies are intended to facilitate reproducible deployment across these environments.

Inference speed remains an important consideration for applications requiring near-real-time processing. On a MacBook Pro equipped with an M4 Pro processor and 24 GB of unified memory, RF-DETR Medium processed 1280×1280 images at approximately 372.90 ms per image, corresponding to approximately 2.68 images per second. This demonstrates practical local inference but does not constitute a direct benchmark of embedded or airborne deployment. Edge deployment on resource-constrained platforms therefore remains an important direction for future work, particularly for applications requiring onboard filtering or real-time feedback (Troscianko et al. 2025; Zábó et al. 2026).

The external results also highlight the importance of confidence calibration and threshold selection. Counting performance was evaluated using a model-specific threshold selected to minimise image-level MAE, meaning that the reported counting results represent an optimised operating point rather than a universal confidence threshold. The Hippopotamus results illustrate why this distinction matters: a threshold that reduces false-negative counting error in densely populated images can simultaneously increase false-positive detections in sparse images. Future work could therefore investigate class- and dataset-specific calibration, alternative counting objectives, or thresholding procedures that explicitly account for spatially heterogeneous target density.

Further improvements may also arise from targeted fine-tuning on underrepresented taxa, improved representation of small objects, and explicit calibration of confidence scores across different flight altitudes and imaging conditions. The training datasets predominantly represent aerial surveys conducted at flight heights of approximately 50–120 m, providing the strongest empirical basis for deployment within this altitude range. We therefore consider 50–120 m to represent the principal operating range supported by the current training distribution. Performance outside this range, particularly at substantially higher altitudes, may be limited by reduced target size and image resolution, as demonstrated by the severe degradation observed on SAVMAP.

More broadly, the present framework provides a general detection layer on which downstream ecological analyses can be constructed. Future work could evaluate the complete pipeline from detection through species classification, tracking, re-identification and behavioural inference, allowing the practical value of bounding-box outputs to be quantified beyond image-level counting.

## 5 Conclusion

Aerial wildlife monitoring requires detection systems that can generalise across taxa, habitats, sensors and survey conditions while remaining useful for more than a single downstream task. Existing specialist counting approaches can provide strong point-based performance, but their outputs are limited primarily to object locations or counts. The results presented here show that a general-purpose RF-DETR detector can retain competitive point-localisation and counting performance while additionally producing explicit bounding-box detections.

On the comparative Cattle dataset, RF-DETR matched OWL across the reported localisation and counting metrics, while on Caribou it achieved almost identical point-based detection performance and similar aggregate counting error. Across the broader external benchmark, specialist systems remained stronger on several datasets, and transfer to unseen avian and hippopotamus taxa was variable. The Hippopotamus results additionally demonstrate that aggregate counting bias can arise from confidence-threshold calibration even when localisation performance remains comparatively strong. At high altitude on SAVMAP, severe performance degradation affected all evaluated architectures, identifying target scale as a shared limitation rather than a detector-specific failure.

Overall, the evaluation provides no evidence of a general trade-off between retaining bounding-box outputs and counting performance. Instead, RF-DETR can approach the broad performance range of specialist wildlife counting systems under several substantially different evaluation conditions while retaining richer spatial information. This combination makes the proposed detector suitable as a general-purpose first-stage detection layer for aerial wildlife monitoring, with counting, classification, tracking and other ecological analyses potentially performed from the same underlying detections.

## 6 Author statements

### 6.1 Acknowledgements

We thank Lancaster University High End Computing (HEC) for providing the GPUs used for model selection training and out-of-distribution dataset evaluation. We would also like to thank Liverpool John Moores University for letting us use the Prospero GPU cluster for the full model training.

### 6.3 Author Contributions

**Chaim Chai Elchik**: Conceptualisation, Manuscript Writing, Code Development, Model Evaluation, Manuscript Writing, Manuscript Editing, Data Annotation.

**Steven Longmore**: Code Development, Manuscript Editing

**Dan Morris**: Model Evaluation, Code Development, Manuscript Editing

**Ben Weinstein**: Model Evaluation, Manuscript Editing

**Josh Veitch-Michaelis**: Model Evaluation, Manuscript Editing

**Camee Knops**: Data Annotation

**Freya Richards**: Data Annotation, Dataset acquisition

**Serge Wich**: Conceptualisation, Manuscript Writing, Data Annotation, Dataset acquisition, Manuscript Editing

### 6.5 Data Availability Statement

The non-public data that support the findings of this study are available from the corresponding author upon reasonable request.

### 6.6 Declaration of generative AI and AI-assisted technologies in the manuscript preparation process

During the preparation of this work, the author(s) used GPT-5 and Gemini Pro 3.1 Extended to assist with writing by rephrasing paragraphs and refining sentence structure, as well as for select code debugging. After using these tools, the author(s) reviewed and edited the content as needed and take full responsibility for the content of the published article.

## 7 Supplementary Materials

**Table 9:** Overview of datasets, source links, and their corresponding licenses.

| Citation | Dataset Link | Dataset Name | License |
| --- | --- | --- | --- |
| Shao et al. 2020 | datasetninja.com/cattle-detection... | DatasetNinja | CC BY 4.0 |
| Adrien Lebreton and Louise Helary 2026 | zenodo.org/records/10400302 |  | CC BY 4.0 |
| L. Helary and A. Lebreton 2024 | zenodo.org/records/11048412 | Cattle_drone_images | CC BY 4.0 |
| Fonod et al. 2025 | zenodo.org/records/13828408 | SONDOGO | CC BY 4.0 |
| Samiappan, B Santhana Krishnan, et al. 2024a | scholarsjunction.msstate.edu/... | AWIR_COW;<br>AWIR_HORSE;<br>AWIR_DEER | CC BY 4.0 (Open Access) |
| sugamg 2026 | github.com/Lmah0/Drone-Wildlife... | WAID_Final | MIT License |
| Delplanque, Linchant, et al. 2024b | dataverse.uliege.be/... | Liege | CC BY 4.0 |
| Mou et al. 2023 | github.com/xiaohuicui/WAID... | WAID_Main | CC BY 4.0 (Knowledge-Sharing) |
| Kline, Kholiavchenko, et al. 2025 | huggingface.co/.../mmla_mpala | MMLA_MPALA | CC BY-NC 4.0 |
| Jenna Kline 2025 | huggingface.co/.../mmla_wilds | MMLA_WILDS | CC BY-NC 4.0 |
| Kline, Nguyen Ngoc, et al. 2025 | huggingface.co/.../mmla_opc | MMLA_OPC | CC BY-NC 4.0 |
| Koger et al. 2023 | edmond.mpg.de/.../3.EMRZGH | Koger | CC BY 4.0 |
| Price, Pranav C Khandelwal, et al. 2025a | darus.uni-stuttgart.de/.../DARUS-5162 | DAZZLE | CC BY 4.0 |
| Naik et al. 2024 | edmond.mpg.de/.../3.JCZ9WK | Bucktales | CC BY 4.0 |
| J. A. Eikelboom et al. 2019 | data.4tu.nl/.../12713903/1 | Eikelboom | CC BY 4.0 |
| Akshatha et al. 2023 | learnermanipal-my.sharepoint.com/... | MIT_Person | Proprietary |
| Kumar et al. 2024 | github.com/UCSB-VRL/WildlifeMapper | MARA | UCSB License |
| Praschl et al. 2026 | github.com/bambi-eco/Dataset | BAMBI | CC BY 4.0 |

